# Balancing Act: A Neural Trade-off Between Coherence and Creativity in Spontaneous Speech

**DOI:** 10.1101/2024.12.23.630053

**Authors:** Tanvi Patel, Sarah E. Macpherson, Paul Hoffman

**Affiliations:** Human Cognitive Neuroscience, Department of Psychology, School of Philosophy, Psychology & Language Sciences, University of Edinburgh, UK

**Keywords:** Discourse, naturalistic speech production, speech coherence, creativity, divergent semantic integration, network analyses

## Abstract

Effective communication involves a delicate balance between generating novel, engaging content and maintaining a coherent narrative. The neural mechanisms underlying this balance between coherence and creativity in discourse production remain unexplored. The aim of the current study was to investigate the relationship between coherence and creativity in spontaneous speech, with a specific focus on the interaction among three key neural networks: the Default Mode Network, Multiple Demand Network, and the Semantic Control Network. To this end, we conducted a two-part analysis. At the behavioural level, we analysed speech samples produced in response to topic cues, computing measures of global coherence (indexing the degree of connectedness to the main topic) and Divergent Semantic Integration (DSI; reflecting the diversity of ideas incorporated in the narrative). Coherence and divergence in speech were negatively correlated, suggesting a trade-off between maintaining a coherent narrative structure and incorporating creative elements. At the neural level, higher global coherence was associated with greater activation in the Multiple Demand Network, emphasising its role in organising and sustaining logical flow in discourse production. In contrast, functional connectivity analyses demonstrated that higher DSI was related to greater coupling between the Default Mode and Multiple Demand Networks, suggesting that creative speech relies on a dynamic interplay between associative and executive processes. These results provide new insights into the cognitive and neural processes underpinning spontaneous speech production, highlighting the complex interplay between different brain networks in managing competing demands of being coherent and creative.

## Introduction

What makes a speaker interesting? Leaving aside the razzle dazzle of gestures, intonation and other non-verbal communication, an interesting speaker distinguishes themselves by delivering content that goes beyond the ordinary. These speakers bring new ideas and provide unconventional perspectives to a given topic. To explore these different angles on a topic, speakers rely on divergent thinking, i.e., the ability to generate multiple different ideas from a single starting point, an ability that underpins creativity in many domains (Guilford, 1967). However, to be effective communicators, they must also ensure that their discourse remains coherent, i.e., that these disparate ideas are organized logically and weaved into a structured and well-connected narrative (Glosser & Deser, 1992). Indeed, a conversational cue may invoke a range of semantic information in the mind of a speaker, and it is up to the skilled speaker to select the subset of information that may be most relevant - and, hopefully, most interesting.

Recent advances in network neuroscience have implicated the Default Mode Network, the Multiple Demand Network (also referred to as the ‘Executive Control Network’ in creativity research), and the Semantic Control Network in both creative processes (Luchini & Beaty, 2023) and in the production of coherent speech (Hoffman, 2019; Morales et al., 2022). However, the interplay between these networks and their specific contributions to creativity and coherence in spontaneous narrative speech production remain largely unexplored. To understand the neurocognitive mechanisms that make a speaker engaging and effective, the present study explored the relationships between creativity and coherence when individuals generate extended passages of speech. To provide the context for this investigation, we first review theories of the neural basis of creativity before considering how these processes relate to discourse production.

### The neural basis of creativity

Creative thought involves a complex interplay between spontaneous, bottom-up associative processes and top-down, controlled executive processes (Kleinmintz et al., 2019). Semantic memory, our stores of knowledge about people, places, and things, plays a central role in the idea generation stage of this process (Abraham, 2014; Abraham & Bubic, 2015). Associative theories suggest that this knowledge is stored as a network of interconnected concepts of varying associative strengths, with the spontaneous spread of activation across this network resulting in the combination of more remote elements (Beaty & Kenett, 2023; Kenett & Faust, 2019; Mednick, 1962). Under this framework, the emphasis is on the individual differences in the structure of semantic memory: creative individuals have many weak associations between concepts, allowing them to bypass stereotypical connections and integrate diverse elements more effectively. Indeed, creativity has been linked to network properties that support the facilitation of remote associations: more creative individuals possess flexible, densely connected networks with shorter path lengths between concepts, enabling activation to spread more efficiently across the network (Kenett, 2018; Kenett et al., 2014, 2018; Kenett & Austerweil, 2016; Kenett & Faust, 2019).

While novelty is an important aspect of creativity, ideas must also be appropriate to the constraints of the task (Runco & Jaeger, 2012). To achieve this balance, during the idea evaluation phase, individuals assess the quality of these ideas by monitoring, evaluating, and selecting them according to the task goals (Kleinmintz et al., 2019). Several top-down processes exert cognitive control over creative processes, including domain-general executive functions like working memory, shifting and inhibition (Benedek et al., 2012; Cheng et al., 2016; Palmiero et al., 2022; Zabelina et al., 2012); fluid and crystallized intelligence (Cho et al., 2010; Forthmann et al., 2019; Nusbaum & Silvia, 2011); and broad retrieval ability (Beaty & Silvia, 2012; Forthmann et al., 2019; Miroshnik et al., 2023). These domain-general control processes facilitate creative thinking by filtering out irrelevant distractors; maintaining and manipulating multiple pieces of information in working memory; and enabling flexible shifting between ideas. Taken together, the literature suggests that creative thinking arises from a dynamic interaction between these spontaneous and controlled processes: ideas are generated through associative processes acting on semantic and episodic memory, and iteratively evaluated for their adherence to the task requirements (Beaty, Silvia, et al., 2014; Kleinmintz et al., 2019).

This dynamic interaction between spontaneous and controlled processes is mirrored in brain activity: creative cognition is mediated by an interaction between brain networks supporting idea generation and goal-directed evaluation (Beaty et al., 2019; Luchini & Beaty, 2023). Resting state functional connectivity and task-based fMRI studies have identified two key networks that dynamically interact to produce novel and appropriate ideas: the Default Mode Network (DMN) and the Multiple Demand Network (MDN) (Adnan, Beaty, Lam, et al., 2019; Adnan, Beaty, Silvia, et al., 2019; Beaty et al., 2015; Shi et al., 2018; Zhu et al., 2017). The DMN consists of a distributed set of regions across the frontal, temporal and parietal cortices, including medial prefrontal cortex (mPFC), posterior cingulate cortex (PCC), precuneus, temporoparietal junction, and the bilateral inferior parietal lobes (IPL; Smallwood et al., 2021). Typically, the DMN shows reduced activation in relation to externally focused tasks, but increases its activation when individuals are engaged in spontaneous and self-generated thought (Andrews-Hanna, 2012; AndrewslJHanna et al., 2014). Consequently, DMN activity has been associated with a variety of internally directed cognitive processes, including mind-wandering (Christoff et al., 2009, 2016; Fox et al., 2015), autobiographical retrieval (Spreng et al., 2009), mental simulation (Hassabis & Maguire, 2007), and episodic future thinking (Cona et al., 2023; Schacter et al., 2012). The DMN has been reported to be active during divergent thinking in several studies (Beaty, Benedek, et al., 2014; Benedek et al., 2014; Ellamil et al., 2012; Fink et al., 2018; Gonen-Yaacovi et al., 2013; Shofty et al., 2022; Takeuchi et al., 2012; Wei et al., 2014). With respect to creativity, the DMN is thought to be involved in the idea generation phase, integrating self-referential, episodic, and semantic information to create novel ideas.

The MDN, consisting primarily of regions in the dorsolateral prefrontal cortex, anterior cingulate cortex, anterior insula and intra-parietal sulcus, is engaged during tasks that require cognitive control (Seeley et al., 2007). This distributed network of regions - also referred to as the ‘Executive Control Network’ (ECN) or ‘Fronto-Parietal Control Network’ (FCPN) (Fedorenko et al., 2013; Yeo et al., 2011) - is involved in top-down control of cognition, including cognitive processes such as working memory, flexibility, and response inhibition (Niendam et al., 2012). Thus, the MDN may be crucial during the idea evaluation phase of creativity, to assess whether candidate ideas meet task criteria and to modify them accordingly (Gonen-Yaacovi et al., 2013). Importantly, creative cognition requires a combination of spontaneous idea generation (mediated by the DMN) and goal directed processing (mediated by the MDN). The two networks generally share an antagonistic relationship, with one network deactivating when the other activates (Fox et al., 2005). However, the networks tend to exhibit functional coupling during tasks involving internally directed processes, including episodic future planning and constructive daydreaming (AndrewslJHanna et al., 2014; Christoff et al., 2009; Gerlach et al., 2014; McMillan et al., 2013; Spreng et al., 2010). Similarly, DMN-MDN coupling has been consistently shown to support creativity across a range of tasks, with the DMN contributing to the generation of potential ideas, while the MDN exerts top-down control to monitor progress in line with task related goals (Beaty et al., 2015, 2016; Green et al., 2015; Liu et al., 2015; Mayseless et al., 2014; Zhu et al., 2017). Indeed, stronger and more efficient connectivity between these regions is associated with individual differences in creativity and better performance on creativity tasks (Beaty et al., 2015, 2018; Green et al., 2015). Moreover, there is increased coupling throughout the creative process, suggesting that creativity benefits from flexible exercise of cognitive control mechanisms (Zabelina & Robinson, 2010).

Aside from the MDN and domain-general executive functions, domain-specific semantic control mechanisms may also have roles in regulating the search, manipulation, and use of task-relevant information in the pursuit of creative goals (CogdelllJBrooke et al., 2020; Krieger-Redwood et al., 2023). Semantic control mechanisms regulate access to and use of semantic knowledge and are associated with a distinct neural network including the left inferior frontal gyrus, posterior middle temporal gyrus and dorsomedial PFC (Jackson, 2021; Noonan et al., 2013; Reilly et al., 2024). The Semantic Control Network (SCN) has been shown to contribute to creativity over and above contributions of DMN and MDN (e.g., Krieger-Redwood et al., 2023; Rastelli et al., 2024). Key areas of the SCN, specifically the left IFG and dorsomedial PFC, are recruited when generating creative links between concepts (Krieger-Redwood et al., 2023) and during the production of creative story narratives (Rastelli et al., 2024), especially when the level of semantic control demands are manipulated. Given the spatial adjacency of the SCN to the DMN and MDN, this may allow for flexible semantic retrieval, i.e., the manipulation of semantic knowledge to fit task relevant goals (Chiou et al., 2023; Wang et al., 2020). The position of the SCN may allow it to leverage interactions between these networks, facilitating the generation of novel and creative ideas by directing attention toward unusual and non-dominant semantic features and associations between concepts.

### Creativity and coherence in discourse

There is now a large literature on the neural and cognitive underpinnings of creativity, but creative thinking is usually assessed with tasks that are rather different from natural discourse settings. Divergent thinking is often operationalised through open-ended tasks that measure an individual’s ability to generate a variety of ideas that are novel and task-appropriate. For example, the Alternate Uses Task (Guilford, 1967) requires participants to list as many novel and creative uses for an everyday object as they can. While effective at evaluating creativity at the word and sentence level (Weinstein et al., 2022), these traditional divergent thinking tasks may not capture the complexity of creative thinking as it occurs in real-world contexts. Everyday creative thinking involves more than simply generating individual ideas. It encompasses the ability to develop these ideas into coherent narratives, solve complex problems, and adapt to diverse situations (Ilha Villanova & Pina E Cunha, 2021). However, work examining these more complex creative outcomes has been limited by the reliance on subjective scoring methods such as expert ratings and rubric-based scoring, that are time consuming, labour intensive and may require expert judges. To address these limitations, advancements in computational linguistic models have enabled automated methods of scoring longer-form responses, which show high correlations with human ratings (Johnson et al., 2022; Orwig et al., 2021). These methods have been used to assess creativity in more complex domains, such as creative writing (D’Souza, 2021; Fan et al., 2023; Zedelius et al., 2019), academic writing (Bower et al., 2023), story generation (Rastelli et al., 2024), and poetry composition (He et al., 2022). Despite these advancements, one area that remains unexplored is spontaneous narrative speech, an important aspect of real-time communication that provides a rich avenue for exploring everyday creativity in an ecologically valid manner.

In the context of narrative speech, being creative may involve generating more divergent content, i.e., ideas that are more distantly connected to the topic under discussion. Additionally, it is crucial for the speaker to combine and organize these ideas in a coherent and engaging manner, ensuring that the overall narrative remains understandable. These dual constraints have been investigated in the domain of creative writing. Human-based scoring of story originality is related to the global semantic distance between the words in a story, suggesting that stories that incorporate a broader array of concepts are judged to be more creative (Fan et al., 2023). At the same time, the global cohesion of the story - the degree to which the story ending was semantically related to the context - is related to human ratings of story rationality, emphasising the importance of building meaningful connections between ideas and to the main story arc. Importantly, higher global cohesion was associated with lower originality, suggesting a trade-off whereby maintaining a coherent and rational structure may limit the inclusion of novel or unconventional ideas. This observed trade-off between coherence and divergence in creative writing underscores the necessity to investigate these dynamics in spontaneous speech.

Just as is seen in creative thinking tasks, the generation of coherent discourse appears to require a complex interplay of associative semantic processes and executive control. Speaking in response to a conversational cue requires both the generation of concepts and ideas (many of which stem from semantic knowledge) and planning and selection processes which ensure that only topic-relevant ideas are included in the discourse and that these are combined in a coherent fashion (Barker et al., 2017; Kintz et al., 2016; Marini & Andreetta, 2016). In line with this view, people with better cognitive control produce more coherent narratives (Hoffman et al., 2018; Kintz et al., 2016; Wright et al., 2014) and, in older people in particular, poor inhibitory function is associated with the production of rambling, verbose narratives (Arbuckle & Gold, 1993). Additionally, recent work highlights a specific role of semantic control processes in speech coherence, over and above domain-general executive abilities (Hoffman et al., 2018). In this study,semantic control abilities positively predicte speech coherence, indicating that the ability to select task-relevant semantic information and suppress irrelevant aspects of knowledge is crucial to coherent communication.

The three networks discussed earlier – the DMN, MDN and SCN – are also implicated in discourse production. The DMN plays a critical role in the construction of high-level situation models that represent the elements of events and experiences and their interrelationships (Yeshurun et al., 2021). These situation models are important for guiding the course of an ongoing narrative. Less coherent discourse places higher demands on this system: DMN activation is higher during the production and comprehension of less coherent discourse (Morales et al., 2022). This increased DMN activation may reflect the regular updating and reconfiguring of mental models required when discourse lacks coherence, suggesting that the DMN is actively engaged in integrating diverse pieces of information and adapting the narrative as it unfolds.

In addition to the DMN, key areas of the SCN have also been implicated in maintaining coherence. When older adults produced passages of extended speech during fMRI, more coherent speech was associated with increased activation in areas of the prefrontal cortex, namely the bilateral IFG (BA45), and the rostro lateral PFC (RLPFC; BA10; Hoffman 2019). Transcranial magnetic stimulation to IFG also impairs narrative coherence (Marini & Urgesi, 2012) and lesions to this region are associated with reduced coherence in post-stroke aphasia (Alyahya et al., 2022). These findings suggest that, while the DMN integrates higher-level semantic information to form situation models, the SCN controls and regulates how this information is incorporated into the unfolding narrative. The SCN’s role may lie in selecting relevant information and inhibiting irrelevant details to maintain a coherent structure. This would be consistent with increased SCN activation under conditions of high semantic competition in a range of other tasks (Badre & Wagner, 2007; Thompson-Schill et al., 1997; Vitello & Rodd, 2015).

Finally, the role of the MDN in coherence is still unclear. Some researchers have proposed that regulation of speech production relies on domain-general executive control regions, particularly those within the dorsolateral PFC (Wise & Geranmayeh, 2016). However, Morales et al., (2022) did not find any modulation of activity in the MDN in relation to coherence during the production of spontaneous naturalistic speech. However, the MDN has been associated with global cohesion in a story generation task (Fan et al., 2023), indicating that the MDN may be significant in relation to tasks that require more structured and deliberate organisation of content.

### The present study

Thus far, we have put forth the argument that, when generating narrative speech, individuals must balance a desire to be divergent (and therefore maximally interesting) with the need to maintain coherence. Evidence from network neuroscience suggests that both these qualities rely on similar underlying neurocognitive networks. On the one hand, creativity is underpinned by a DMN-MDN coupling that allows for an iterative interplay between bottom-up associative processes and top-down executive control during idea generation and evaluation. This coupling is supported by the SCN, enabling flexible, goal-directed semantic retrieval and selection processes. On the other hand, idea generation in discourse is underpinned by DMN-related activity that contributes to building and monitoring of situation models. In addition, both executive functions and semantic control are related to coherence at a behavioural level, and greater SCN activation has been linked with more coherent speech production. However, the neural correlates of creativity and coherence in language generation have previously been studied separately. This is potentially problematic as these qualities are likely to be inter-correlated, with more coherent speech being less creative and vice-versa. With all this in mind, the present study has two aims: (1) to explore how creativity and coherence in language production are related at the behavioural level; (2) to use fMRI to understand how network-level activation and functional connectivity relates to these two constructs.

## Study 1: Methods

In Study 1, we collated a large set of verbal responses to topic prompts from previously published and unpublished data (see below). Then we computed established measures of coherence and creativity (divergence) for all responses, using computational linguistics methods, to examine the relationship between these constructs.

### Participants

For the current sample, we aggregated data from four previous studies: ’*MD Dataset*’ (Unpublished); ’*18-E Life Dataset*’ (Hoffman et al., 2018); ’*19-NC Dataset*’ (Hoffman, 2019); and *SL-19 Dataset* (Morales et al., 2022; Patel et al., 2023; Wu, Morales, et al., 2022). All data were collected at the University of Edinburgh, School of Philosophy, Psychology and Language Sciences. The studies included young adult participants from the student population as well as older adults from the Psychology Department Volunteer Panel. Combining these datasets resulted in a final sample of 153 participants (102F, 51M), of which there were 85 younger (M_Age_= 21.08, SD_Age_ = 3.32, range 18-35, 65F, 20M) and 68 older (M_Age_= 75.53, SD_Age_ = 7.75, range = 60-92, 37F, 31M) adults. For further details on the demographic summary for each study, see Table S1 in the Supplementary Materials. All participants were native English speakers, with no history of neurological or psychiatric illness. The studies received ethical approval from the Edinburgh Psychology Research Ethics Committee, and informed consent was collected from all participants.

### Design and Procedure

In each of the studies, participants were given a series of prompts and asked to generate speech for a period of between 50 and 60s. The prompts were constructed to probe specific aspects of semantic knowledge (e.g., ‘*What would it have been like to live in the Middle Ages*?’). Further details on the procedure adopted in each study, as well as a full list of prompts, are given in the Supplementary Materials.

### Measures

Spoken responses to each prompt were digitally recorded, transcribed, and cleaned. We computed measures of global coherence, divergence (DSI), and the average number of words per response.

#### Global coherence

To quantify the degree to which utterances were related to the topic participants were asked about, we used a computational linguistic measure of global coherence (Hoffman et al., 2018). This measure is based on Latent Semantic Analysis (LSA; Landauer & Dumais, 1997), a technique that uses patterns of word co-occurrence to build a high-dimensional semantic space. A co-occurrence matrix, generated from a large corpus of natural language, is used to encode the frequency of words in different contexts. Singular value decomposition is used to reduce the matrix dimensionality such that each word is represented as a high-dimensional vector (or embedding), with words used in similar contexts allotted similar vectors. Following Hoffman et al. (2018), for each speech sample, we used LSA to generate a vector-based representation of its semantic content. Next, we computed a composite vector by averaging the LSA representations of all participants’ responses to the target prompt (excluding the target response itself). This composite vector was taken to represent a prototypical response to the prompt. Finally, for the target response, we constructed smaller windows of 20 words each, containing the current word and 19 preceding words. For every 20-word window, we computed an LSA vector to represent the semantic content. Global coherence was assessed by measuring the cosine similarity of the vector for each window with the vector representing a typical response to the prompt. Higher global coherence values represented discourse more typical of the topic, while lower coherence values indicated less topic-relevant speech. These values were averaged to provide a mean global coherence for each response. Global coherence calculated in this way has been shown to have high test-retest reliability and to be strongly related to human judgements of coherence (Hoffman et al., 2018). Global coherence values can theoretically range from 0 to 1, but the range observed in previous studies has been from .20 to .80.

#### Divergent semantic integration

To assess the degree to which the responses incorporate divergent ideas, we used DSI, an established measured of semantic diversity for discourse samples (Johnson et al., 2022). DSI indexes the semantic distance between words in a given response, using context-dependent word vectors from a large language model - bidirectional encoder representations from transformers (BERT; Devlin et al., 2019). BERT produces context-dependent vectors (embeddings) that represent the semantics of each word in a sample, taking into account the surrounding linguistic context. These embeddings are then used to calculate DSI: the embeddings for each word in a sentence are compared to each other using pairwise cosine semantic distance, and these are averaged across all word pairs in the response. This aggregate distance provides a single DSI score for the response, representing the degree to which the response integrates more diverse semantic information. High DSI scores suggest that the response integrates more divergent ideas, while low scores suggest less semantic diversity. DSI has been shown to be a good predictor of creativity in narrative text, showing strong correlations with human ratings of creativity (Johnson et al., 2022; Orwig et al., 2023). DSI values can theoretically range from 0 to 1, but the range observed in previous studies has been from .70 to .90 (Orwig et al., 2024).

#### Number of words

We computed the number of words in each response, as length of the response is potentially correlated with both DSI and coherence (Johnson et al., 2022).

### Analysis Strategy

First, we used Pearson correlational analysis to understand the relationship between global coherence and DSI, and their relationship with our covariates of interest, i.e., the number of words produced and age. To further explore this relationship, we used a linear mixed effects model to examine whether coherence predicts DSI, after controlling for age and the number of words in the response. We fitted the model with maximum likelihood estimation and the bobyqa optimiser, using the lme4 package (Bates et al., 2015). We included random intercepts for participants nested within the datasets, to account for variation across participants in the different datasets, and random by-prompt intercepts, to account for variation across the different prompts. Sum to zero coding was adopted for age group, with younger adults as the first level.

## Study 1: Results

### Descriptive and correlational analysis of speech metrics

Over the 5 datasets, participants generated 2057 responses to 28 distinct prompts (for a full list of prompts, see Supplementary Materials). We excluded 9 responses that were less than 50 words long, leaving us with a final sample of 2048 responses. Descriptive statistics for the measures of interest are presented in Table 1. Overall, participants produced an average of 133.43 words per response. Independent sample t-tests indicated that, compared to their older counterparts, younger people produced speech that was less divergent (*t*(150.99) = -3.17, *p* <.01) and more coherent (*t*(142.01) = 3.58, *p* <.001; see Figure 1a and 1b). This supports previous work suggesting that older people tend to be less coherent and produce speech that contains more semantically diverse topics (e.g., Hoffman et al., 2018; Wright et al., 2014).

**Figure 1:**
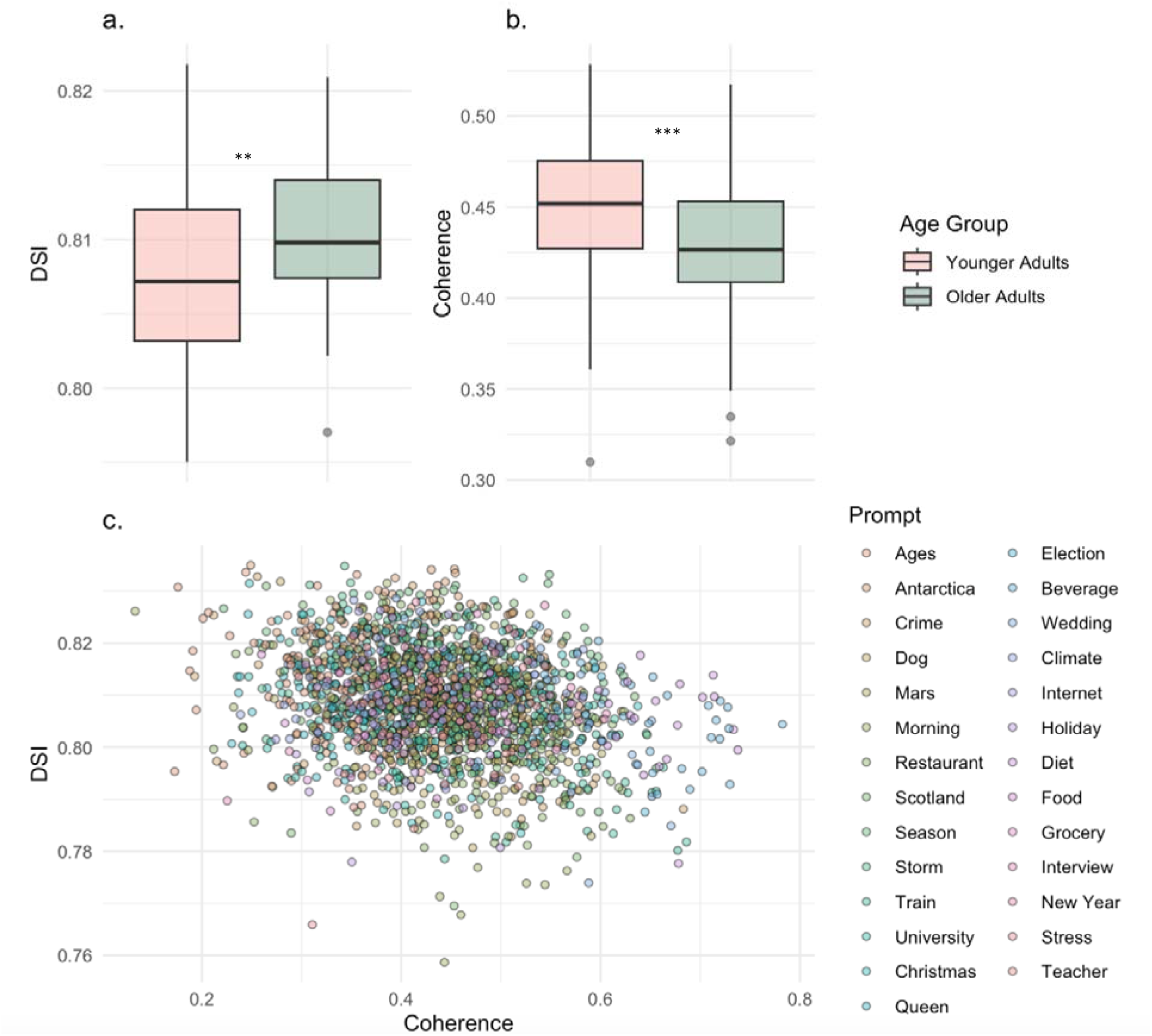
DSI and Coherence Across Age Groups and Prompts. (a) Boxplot of DSI scores in younger and older people; (b) Boxplot of Coherence scores in younger and older people. (c) Scatterplot showing the relationship between DSI and Coherence scores across experimental prompts. *\*\* p &<.01, *** p &<.001*.

**Table 1:** Descriptive Statistics for Measures Across Datasets and Overall.

| Measure | Group | Mean | SD | Min | Max |
| --- | --- | --- | --- | --- | --- |
| Coherence | MD | 0.43 | 0.08 | 0.13 | 0.63 |
|  | 18-ELife | 0.44 | 0.09 | 0.20 | 0.69 |
|  | 19-NC | 0.45 | 0.11 | 0.19 | 0.78 |
|  | 19-SL | 0.47 | 0.10 | 0.23 | 0.74 |
|  | **Overall** | **0.44** | **0.09** | **0.13** | **0.78** |
| DSI | MD | 0.81 | 0.01 | 0.77 | 0.83 |
|  | 18-ELife | 0.807 | 0.011 | 0.759 | 0.835 |
|  | 19-NC | 0.807 | 0.010 | 0.774 | 0.832 |
|  | 19-SL | 0.808 | 0.009 | 0.766 | 0.829 |
|  | **Overall** | **0.81** | **0.01** | **0.76** | **0.84** |
| No of Words | MD | 146.28 | 36.83 | 53 | 281 |
|  | 18-ELife | 138.03 | 34.97 | 50 | 244 |
|  | 19-NC | 103.18 | 18.50 | 50 | 152 |
|  | 19-SL | 124.32 | 25.14 | 67 | 196 |
|  | **Overall** | **133.43** | **35.38** | **50** | **281** |

To examine the associations between the variables, we computed Pearson correlation coefficients at the response-level between the measures of interest in each individual dataset and in the combined dataset (see Table 2). In the combined dataset, coherence was negatively correlated with DSI (*r* = -.23, *p* <.001), indicating a possible trade-off between coherence and divergence. This negative association between global coherence and DSI persisted in the individual datasets, although this association did not consistently reach significance. The number of words generated in the response was negatively related to coherence (*r* = -.07, *p* <.01).

**Table 2:** Correlations between response-level measures across individual samples and combined dataset. *Note: DSI = Divergent Semantic Integration*.

| Group | Coherence - DSI |  | Coherence – Words |  | DSI – Words |  |
| --- | --- | --- | --- | --- | --- | --- |
|  | *r* | *p* | *R* | *p* | *r* | *p* |
| MD | -.25 | <.001 | -.13 | <.001 | .20 | <.001 |
| 18-ELife | -.30 | <.001 | .007 | .83 | -.17 | <.001 |
| 19-NC | -.14 | <.05 | .18 | <.01 | -.006 | .90 |
| 19-SL | -.08 | .13 | -.08 | .14 | -.03 | .52 |
| Overall | -.23 | <.001 | -.07 | <.01 | .04 | .09 |

### Global Coherence as a Predictor of Semantic Divergence in Speech

We used a mixed effects model to test whether global coherence predicted DSI after accounting for confounds, i.e., number of words and age group. All variables were scaled prior to inclusion in the model. Coherence had a significant negative effect on DSI, indicating that increases in global coherence were associated with a decrease in DSI (B = -0.13, 95% CI [-0.18, -0.09], *p* <0.001). Moreover, age was a significant predictor, with younger individuals demonstrating lower DSI scores overall when compared to older adults (B = -0.33, 95% CI [- 0.44, -0.21], *p* <0.001). The number of words generated was not a significant predictor of DSI (B = -0.01, 95% CI [-0.07, .04], *p* =0.67). However, the pattern of effects was present in each age group, suggesting that the negative relationship between DSI and coherence holds across different ages, although younger individuals overall tend to have lower DSI scores.

## Study 1: Discussion

In Study 1, we examined whether coherence of discourse is related to its creative quality, measured using the Divergent Semantic Index (DSI). Our analysis of 2048 responses across five datasets revealed a significant negative association between coherence and DSI, suggesting that coherence and divergence in narrative speech are inherently linked, with higher coherence generally leading to lower divergence and vice versa. This trade-off highlights the complex balance individuals must maintain between generating divergent ideas and producing structured, on-topic speech. It also highlights the importance of controlling for divergence when assessing effects of coherence and vice-versa. Building on these insights, Study 2 aimed to investigate the neural mechanisms underpinning the observed trade-off between coherence and divergence. Using activation and functional connectivity analyses, we investigated how different brain networks interact to support creative and coherent speech. Specifically, we examined the roles of the Default Mode Network (DMN), the Multiple Demand Network (MDN) and the Semantic Control Network (SCN) in mediating the balance between creative divergence and coherence in discourse.

## Study 2: Methods

Study 2 reports analyses of the 19-SL fMRI dataset, in which participants produced 50 second discourse samples in responses to topic prompts. Other analyses of these data have been reported previously (see Morales et al., 2022; Patel et al., 2023), but this is the first study to examine the effects of Divergent Semantic Integration.

### Participants

Twenty-five participants (M_Age_ = 24, SD_Age_ = 4.4, range = 18-35, 21F, 4M) were recruited from the University f Edinburgh student population and compensated for their time. Participants were right-handed (based on the Edinburgh Handedness Inventory; Oldfield, 1971); native English speakers (defined as learning English before age 5); and in good health, with no history of neurological or psychiatric illness. Ethical approval was received from the Edinburgh Psychology Research Ethics Committee and all participants gave informed consent.

### Design and Procedure

Participants were required to either generate or listen to naturalistic speech in 50 second blocks while in the scanner. The experiment consisted of 4 runs in total, with each participant completing two runs of production and two runs of comprehension. The order of the runs was counterbalanced across participants, and each run lasted approximately 8 and a half minutes. For every run, there were 6 discourse trials and 5 baseline trials, randomised for each participant. The current study only analysed the speech production runs: for each trial, participants were presented with the prompt on a screen for 6s, to allow them to prepare their speech. They were instructed to start speaking when a green circle appeared in the centre of the screen, and to stop when a red cross appeared at the end of the 50s period. In the baseline condition, participants recited the popular British nursery rhyme ‘Humpty Dumpty’ for 10s and were asked to start again from the beginning if they finished early (see Figure 2a). The interstimulus interval between trials was jittered between 3-7 seconds (M = 5 seconds). The stimuli were presented using E-prime software on a screen behind the scanner, viewed on a mirror placed on the head coil. Participants were given two practice trials at the start to familiarise themselves with the production task and the baseline condition

To elicit speech production, we devised 12 prompts related to common semantic knowledge on a series of topics (e.g., ‘*Do you think the internet has improved people’s lives?’* See Supplementary Table S3 for a list of all prompts used). These semantic speech conditions were contrasted with a baseline condition, in which participants recited the nursery rhyme. This active low-level baseline was chosen as it consists of grammatically sound yet automatic speech that requires minimal semantic processing.

### Processing of speech samples

The stages in the analysis pipeline are illustrated in Figure 2b. Speech was recorded using an MRI-compatible microphone, processed with noise cancellation software to reduce scanner noise (Cusack et al., 2005) and transcribed for analysis.

#### Global coherence

To index global coherence, we used the same computational linguistic measure as Study 1. A global coherence score was calculated for each response and used as a predictor of BOLD responses and connectivity (Hoffman, 2019).

#### Divergent semantic integration

As in Study 1, we used DSI to index the degree to which each response connected divergent ideas (Johnson et al., 2022).

#### Number of words

We computed the number of words in each response, as the length of the response may be a potential covariate affecting DSI scores (Johnson et al., 2022) and global coherence.

### Image Acquisition and Processing

Participants were scanned in a 3T Siemens Prisma Scanner using a 32-channel head coil. Functional image data were acquired at three echo times (13ms, 31ms and 48ms) using a whole brain multiband multi-echo acquisition protocol (Feinberg et al., 2010; Moeller et al., 2010; Xu et al., 2013). Pre-processing was carried out using Statistical Parametric Mapping 12 (SPM 12, v7219) and TE-Dependent Analysis Toolbox 0.0.7 (Tedana; DuPre et al., 2019). The three-echo series were weighted and combined into a single time series, and denoised using Tedana. By partitioning the data into BOLD and non-BOLD components, this approach can improve signal quality, remove head movement-related artefacts and reduce susceptibility artifacts commonly seen in orbitofrontal regions and the ventral Anterior Temporal Lobes (ATL; Kundu et al., 2017). Forty-six image slices covering the whole brain were obtained with a TR = 1.7s, 80 x 80 matrix, flip angle = 73°, and isotropic voxel size of 3mm. Over each run, 281 volumes were acquired. For normalisation, we also acquired T1-weighted structural images using an MP-RAGE sequence with 1mm isotropic voxels, TR = 2.5 s, TE = 4.6 ms, flip angle = 7°, in-plane resolution = 224 × 304 matrix, 0.8mm x 0.8mm; 288 slices of slice thickness 0.8mm were acquired (no slice gap). Pre-processed images were then unwarped and spatially normalised to Montreal Neurological Institute (MNI) space using SPM Dartel (Ashburner 2007), which allows for precise inter-subject registration and increased analytic sensitivity. These spatially normalised images were then smoothed using a Gaussian kernel with 8mm full width at half maximum.

**Figure 2.**
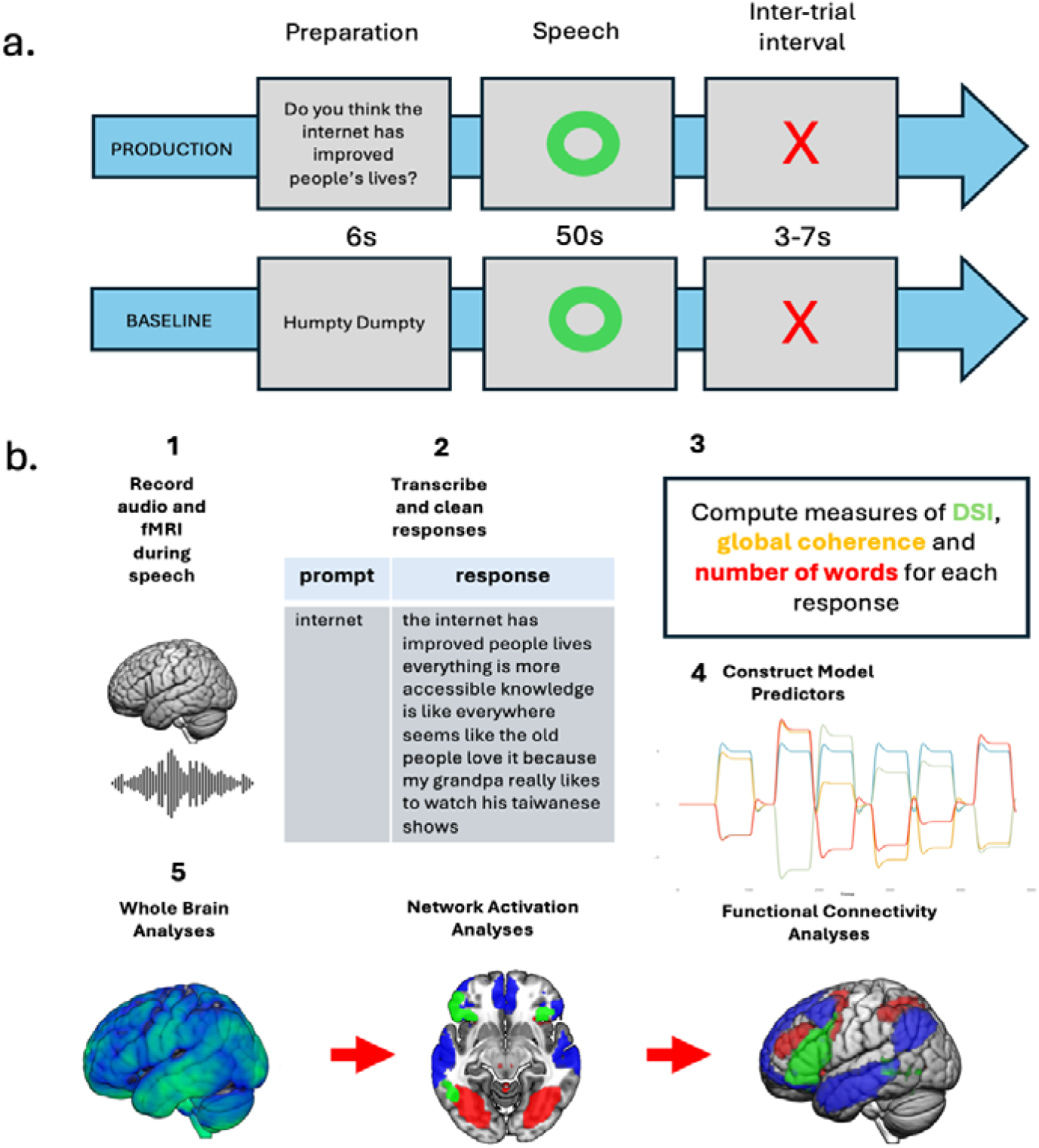
(a) Illustration of a production/baseline trial. (b) Workflow from data collection to neural analysis from (i) Recording of audio and fMRI activity during production; (ii) Transcription; (iii) Computation of coherence, divergent semantic integration (DSI) and number of words; (iv) Constructing model predictors; (v) Stages in Neural Analysis.

### Analyses

#### Participant-level models

First, we estimated participant-level models to get individual-level beta maps representing the effects of discourse characteristics on activation. Four event types were modelled for each run: discourse prompts and baseline prompts (6s duration); discourse production (50s duration) and baseline speech production (10s duration). For the discourse production events, 3 mean-centred parametric modulators were used, representing the global coherence, DSI and number of words for each response. Additionally, 6 motion parameters and their first derivatives were added to the model as covariates. Event regressors were convolved with the haemodynamic response function using SPM12 and data were treated with a high-pass filter with a cut-off of 128s. To assess the brain regions involved in discourse production, we contrasted brain activation for discourse against the baseline task. To assess effects of discourse characteristics on activation, we used the three parametric modulators of the discourse effect.

#### Whole-brain level

First, we investigated whole brain activation at the group-level. For the discourse vs baseline speech contrast, we used a voxel-level threshold of p <0.001 with correction for multiple comparisons (p >0.05) at the cluster level using random field theory. For the effects of global coherence, DSI, and number of words, few regions were significant at this stringent threshold. Therefore, to increase power, our main analyses of these effects focused on 3 large-scale brain networks, described in the following sections. Unthresholded effect maps have been included in the Supplementary Materials (Figure S1) to visualise these effects across the whole brain.

#### Networks of Interest

Our analyses were performed on three networks of interest that have been shown to be related to both coherence and creativity. The networks and masks used to identify them were the same as those used in (Morales et al., 2022):

1. The Default Mode Network (DMN) was identified from a 7 network parcellation of resting-state fMRI data from 1000 participants reported by Yeo et al., (2011). The DMN is shown in blue in Figure 4a, and includes regions of the medial prefrontal cortex, posterior cingulate cortex, anterior temporal lobe, precuneus, and lateral parietal cortex.
2. The Semantic Control Network (SCN) was defined using significant voxels identified in a meta-analysis by Jackson (2021), which highlights regions consistently associated with high cognitive control demands in semantic processing tasks. SCN regions are shown in green in Figure 4a, and include the left inferior frontal gyrus, the left posterior middle temporal gyrus, and presupplementary motor regions.
3. The Multiple-Demand Network (MDN) was defined as the regions associated with increased cognitive demands across multiple domains, as reported by (Fedorenko et al., 2013). The MD network is shown in red in Figure 4a, and includes regions in the dorsolateral PFC, precentral gyrus, insular cortex, intraparietal sulcus, presupplementary and supplementary motor areas, anterior and middle cingulate cortex.

#### Identification of networks using Independent Components Analysis (ICA)

The main aim of this analysis was to identify the effects of discourse coherence, DSI and the number of words on activation in our three networks of interest. To extract the BOLD timecourses in our networks of interest, we used Independent Components Analysis (ICA), a data reduction technique that separates fMRI signals into spatially or temporally independent components that correspond to different brain networks (Calhoun et al., 2009). We used a spatially-constrained semi-blind version of ICA, which extracts independent components from the data that align with pre-specified spatial templates (Lin et al., 2010). This was implemented in Matlab using the Group ICA of fMRI toolbox (GIFT; https://trendscenter.org/software/gift/). We ran the spatially-constrained ICA separately on each participant’s data, using our pre-determined network masks, to constrain the extraction of components. This ensured we recovered exactly three components from each participant’s data, corresponding to our networks of interest.

#### Activation Analyses

Next, we ran a series of general linear models to investigate the effects of coherence and DSI on activation in each network using the BOLD timecourses extracted from the GIFT ICA analysis. Data were treated with a high pass filter with a cut-off of 180s. The regressors in the GLM were identical to those used in the whole-brain analyses, but here, they were used to predict the BOLD response in each network, rather than the response in individual voxels. Contrast values representing the difference in activation between semantic and baseline speech and the effects of GC, DSI and the number of words were computed for each network. Data were aggregated by averaging over the two production runs. T-tests (FDR-corrected) were conducted on the effects in each network to determine whether these were significantly different from zero. We tested whether effects differed between networks using one-way repeated measures ANOVAs, with network type (DMN, MDN, SCN) as the within-subjects factor, and participants as the random effect. Sum to zero coding was used for network type. Following this, pairwise comparisons using paired t-tests with Holm adjustment were used to compare networks.

#### Functional connectivity analyses

Finally, we explored how the functional connectivity between our networks of interest were influenced by the coherence and creativity of the responses participants made. Psychophysiological Interactions (PPI) is a popular type of functional connectivity analysis, which evaluates changes in connectivity strength induced by specific cognitive tasks or conditions. We used the generalised psychophysiological interactions method (gPPI; McLaren, Ries, Xu, & Johnson, 2012) that allows for multiple PPI effects within a single general linear model. The gPPI method examines how the functional coupling between a seed region and other brain regions change as a function of experimental conditions. We used this method to understand how the strength of the correlations between our three large-scale networks of interest varied as a function of our cognitive predictors.

While traditional PPI analyses use small, often spherical, seed regions, we were interested in functional interactions between large-scale networks. For our seeds, we therefore used the BOLD timecourse data extracted for each network of interest (Goulden et al., 2014; Masson et al., 2020). We used the gPPI toolbox to generate task, seed, and PPI regressors for our models using each network as a seed. We then used these models to predict the timecourses in a target network, e.g., DMN-seeded models were used to predict the timecourses of the SCN and MDN. These analyses demonstrated how the between-network correlations were influenced by discourse factors. PPI effects are considered to be bi-directional, so effects from network pairs (e.g., DMN-to-SCN and SCN-to-DMN) were averaged.

We tested the effects of our predictors on the functional connectivity between the ICA-derived networks. First, one-sample t-tests (FDR-corrected) were used to determine whether each discourse factor (discourse vs. baseline, coherence, DSI and number of words) influenced functional connectivity between each network pair. Next, we ran repeated measures one-way ANOVAs, with network pairs as the sum-to-zero coded, within-subject variable and participants as the random effect, to determine whether effects varied between network pairs. Pairwise comparisons using paired t-tests with Holm adjustment were then conducted to compare network pairs.

## Study 2: Results

### Whole Brain Analyses

First, we examined brain activation for discourse versus baseline speech (Figure 3). The production of discourse activated areas typically associated with semantic processing, including the bilateral anterior temporal cortices, superior and inferior frontal gyrus (BA 45 and 47) and the angular gyrus (Binder & Desai, 2011). Activation was seen across regions of the SCN and DMN.

**Figure 3:**
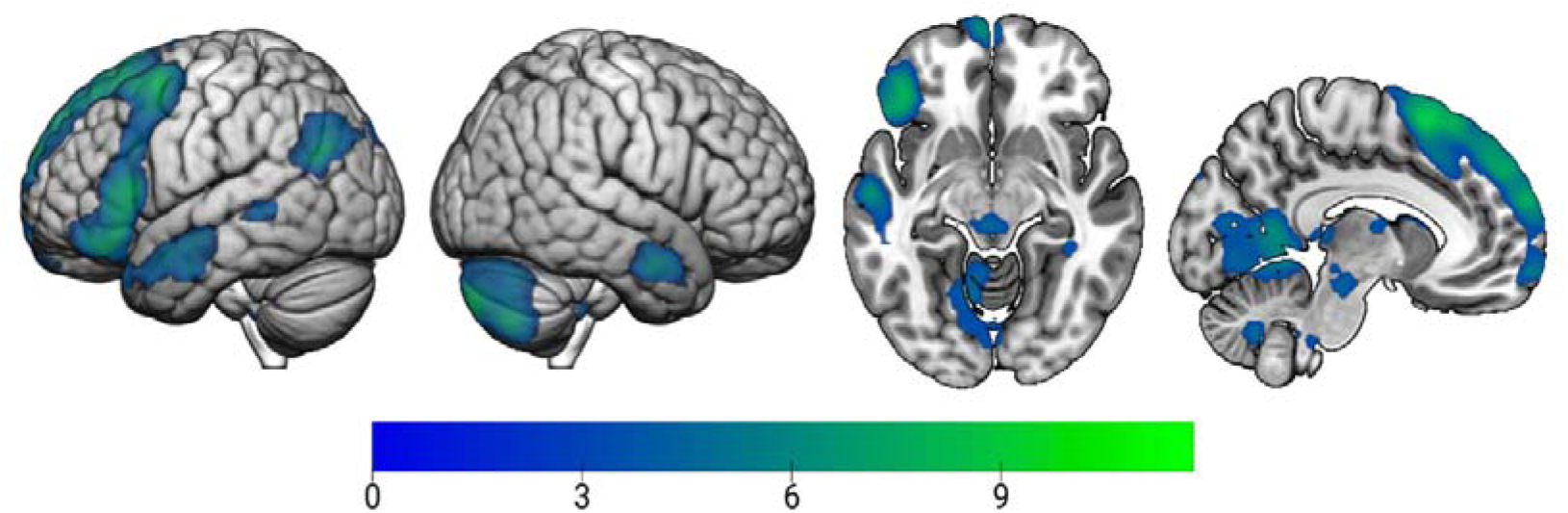
Whole brain activation for discourse compared to baseline speech. Note: Images thresholded at cluster-corrected p&<.05.

### Network Activation Analyses

The DMN and SCN showed increased activation during discourse production, while the MDN showed more activation in the baseline condition, as shown in Figure 4b. FDR-corrected one sample t-tests indicated that all three contrasts were significantly different from zero (see Table 3). The ANOVA results demonstrated a significant main effect of network, F(2, 46) = 88.87, *p* <0.001. All pairwise comparisons were statistically significant, indicating that each network had significantly different contrast values to the other (adjusted *p* <0.001).

Figure 4b shows the effects of different discourse characteristics on activation in each network. Positive effects indicate that higher values of the property were associated with greater levels of activation. The MDN and SCN both showed increased activation when people produced more coherent speech, while the DMN showed increased activation for less coherent speech. The effect in the MDN was highly significant while weaker effects in the SCN and DMN did not survive corrections for multiple comparisons (see Table 3). Further, the repeated measures one-way ANOVA revealed a significant effect of network type (F (2,46) = 17.32, *p* <0.001). Pairwise comparisons indicated significant differences between the DMN and both the MDN and SCN (p>0.001), but not between the MDN and SCN (adjusted *p* =0.34). The significant difference between the DMN, with the MDN and with the SCN, suggests that this network plays a distinct role in relation to maintaining speech coherence. The MDN and SCN did not differ significantly from each other, indicating there may be some shared functional similarities in their roles. In line with this view, the DMN showed decreased activation while people were being more coherent, while the MDN and SCN showed increased activation when people were being coherent.

With regard to DSI, effects did not significantly differ from zero in any of our networks of interest. Further, the effects of DSI did not differ significantly between networks, F(2,46) = .27, *p* =0.72. Similarly, no significant effects were found for the number of words, F(2,46) = .06, *p* =0.94. This indicates that there were no significant differences in effects across our networks of interest in relation to DSI and words in the response.

**Table 3:**
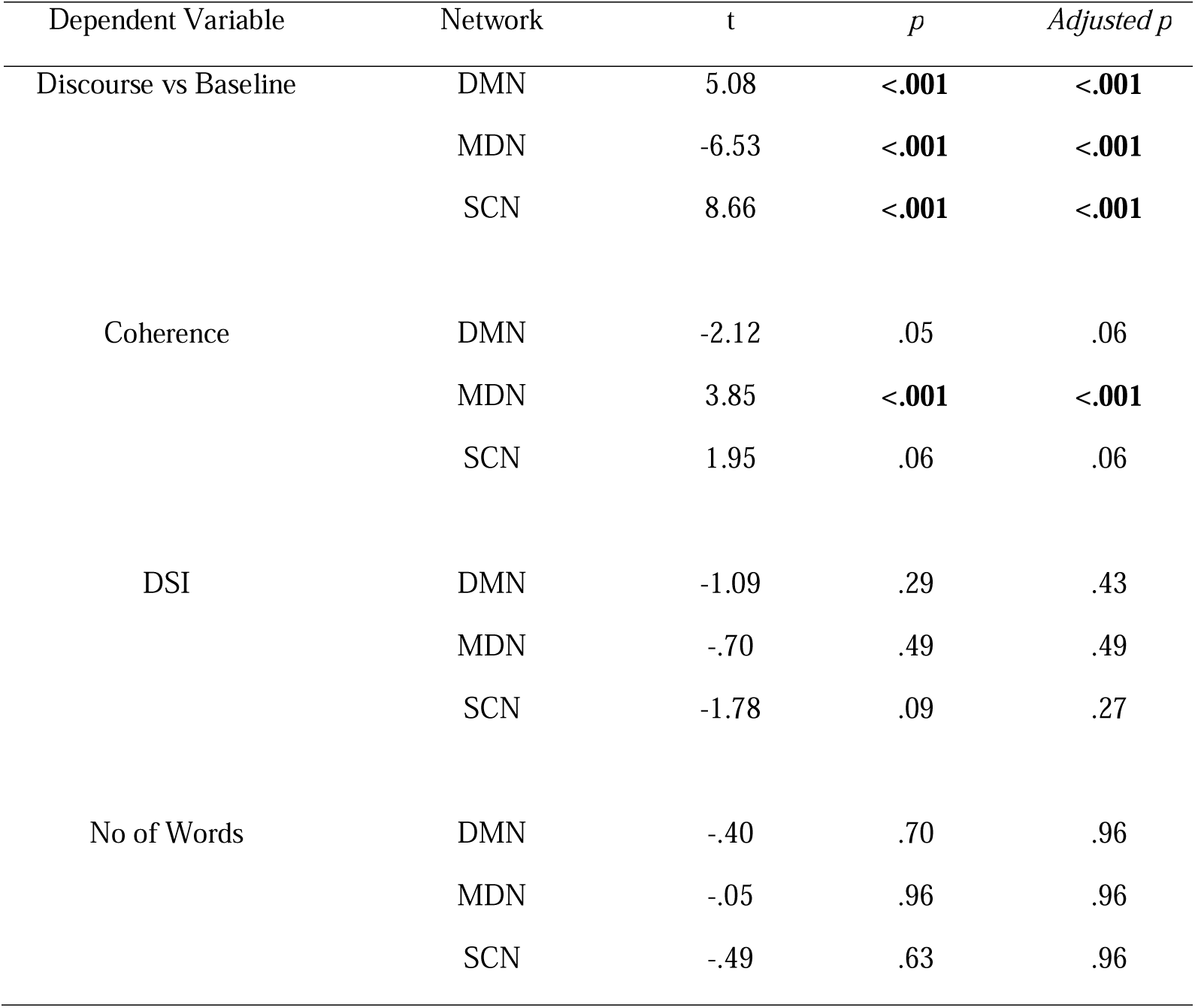
One-sample t-tests for effects across each ICA-derived network.

### Functional Connectivity Analyses

Effects of discourse characteristics on between-network connectivity are presented in Figure 4c. Here, positive effects indicate that higher values of the property were associated with stronger correlations in activity between networks. No network pairs showed connectivity differences in discourse production and baseline speech production (one-sample t-test results shown in Table 4). The ANOVA further confirmed these effects did not differ between our networks of interest, F(2, 46) = 1.06, *p* =0.35. Similarly, between-network connectivity was not associated with coherence and an ANOVA revealed no significant differences in the coherence effects between network pairs, F(2,46) = 2.00, *p* =0.14. In contrast, DSI was associated with changes in connectivity between the DMN and MDN. Connectivity between these two networks was higher when people produced more divergent responses (Figure 4c). Further, an ANOVA suggested that this effect differed from the other two network pairs, F(2,46) = 3.73, *p* =0.03 (although differences were not significant in pairwise comparisons). Finally, the number of words was not associated with changes in connectivity and an ANOVA indicated no differences between network pairs F(2,46) = .85, *p* =0.43.

**Table 4:** One sample t-tests for effects in each network pair.

| Dependent Variable | Network | t | *p* | *Adjusted p* |
| --- | --- | --- | --- | --- |
| Discourse vs Baseline | DMN-MDN | .26 | .80 | .80 |
|  | DMN-SCN | -.93 | .36 | .54 |
|  | MDN-SCN | -1.90 | .07 | .21 |
| Coherence | DMN-MDN | .03 | .98 | .98 |
|  | DMN-SCN | -1.05 | .30 | .46 |
|  | MDN-SCN | 1.12 | .28 | .46 |
| DSI | DMN-MDN | 3.53 | **<.01** | **<.01** |
|  | DMN-SCN | -.02 | .98 | .99 |
|  | MDN-SCN | -.01 | .99 | .99 |
| No of Words | DMN-MDN | 1.64 | .12 | .17 |
|  | DMN-SCN | 2.35 | .03 | .08 |
|  | MDN-SCN | .41 | .68 | .68 |

**Figure 4:**
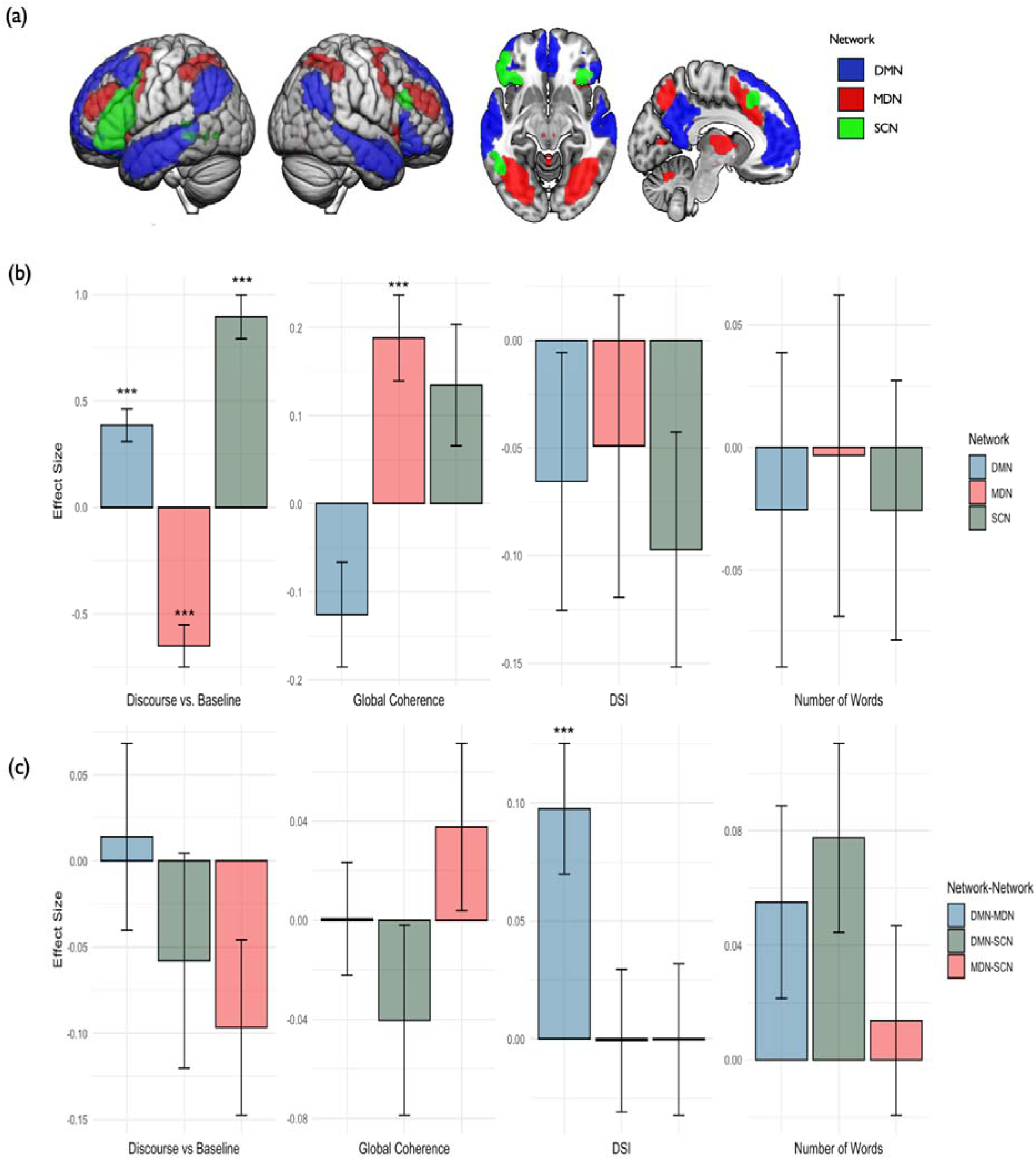
(a) Masks of Networks of Interest; (b) Effects of regressors (semantic vs. baseline speech, global coherence, Divergent Semantic Integration (DSI) and number of words) on network activation; (c) Effects of regressors on functional connectivity between networks. *Note: DMN = Default Mode Network; MDN = Multiple Demand Network; SCN = Semantic Control Network. ** p &<.01, *** p &<.001*

## General Discussion

The aim of this work was to explore the dynamic relationship between coherence and creativity in discourse production. Specifically, we sought to understand the neural activity that gives rise to highly focused vs. more original speech during unscripted verbal communication. To this end, we conducted two studies examining this relationship at the behavioural and neural levels. In the first study, we collated a large set of verbal responses to topic prompts and used established methods to compute measures of global coherence and creativity (Divergent Semantic Integration). We found that the two constructs are negatively related: higher coherence was related to less divergence in speech, suggesting that there is a trade-off between being coherent and creative. In the second study, we examined the neural mechanisms underlying these two constructs. Network activation analyses indicated that more coherent speech was related to increased activation in the MDN. In contrast, functional connectivity analyses suggested that divergence in speech was underpinned by an increased coupling between the DMN and MDN. Together, these findings highlight a complex interplay between these two constructs in discourse. To our knowledge, this is the first study to investigate creativity as an emergent, everyday quality within spoken discourse, a context that has not been explored in traditional creativity assessments.

First, our behavioural study identified a negative relationship between global coherence and divergence in extemporaneous speech. This suggests that when speakers prioritise maintaining a coherent, focused narrative, it may come at the expense of the limiting the originality and diversity of ideas in their speech. In other words, coherence and creativity may function as two opposing forces in communication, where enhancing one may naturally diminish the other. This trade-off is echoed in previous work examining creative story writing: stories that had greater global cohesion, reflecting the degree to which ideas were connected to the central narrative, were rated as being less original (Fan et al., 2023).

The optimal balance between coherence and divergence may differ depending on the context and purpose of communication. Greater coherence may be helpful in contexts where one needs to communicate ideas efficiently and accurately, where any divergence detracts from the main message. However, if the goal of the conversation is to entertain one’s interlocuter, a more meandering conversational style that includes irrelevant details, off-topic comments or unexpected twists could make the interaction more stimulating. Indeed, this trade-off impacts how the audience perceives the speaker, with listeners rating less coherent speakers as being more interesting (James et al. 1998).

Our behavioural results indicated that older people tended to be less coherent and more divergent in their speech, replicating previous findings that older people are more likely to produce off-topic speech (Arbuckle & Gold, 1993; Glosser & Deser, 1992; Hoffman et al., 2018; Kemper et al., 2010; Kintz et al., 2016; Marini et al., 2005; Marini & Andreetta, 2016; Wright et al., 2014). This is thought to stem from age-related declines in domain-general cognitive control mechanisms since more coherent speech is associated with superior attentional and executive control abilities (Arbuckle & Gold, 1993; Hoffman et al., 2018; Kemper et al., 2010; Kintz et al., 2016; Marini & Andreetta, 2016; Wright et al., 2014b). To produce meaningful utterances, one must generate relevant statements while continuously monitoring the flow of ideas. Thus, top-down control is required to select topic-relevant ideas and maintain attentional focus on the subject of interest.

In line with behavioural evidence for the importance of cognitive control in maintaining coherence, our network activation analyses indicated greater speech coherence was related to increased activation in the Multiple Demand Network. This system has been implicated in higher-level cognitive processes that organise, manage, and execute sub-tasks to support the accomplishment of a broader objective (Duncan, 2010; Fedorenko et al., 2013). MDN may optimise speech coherence through planning and regulatory operations such as: planning the sequence of ideas and transitions between them, suppressing irrelevant thoughts that could lead to off-topic speech, monitoring speech for errors, and refocusing speech back to the topic if deviations are detected. Indeed, beyond classical language areas, speech production has been linked to a network of regions associated with domain-general cognitive control (AbdulSabur et al., 2014; Adank, 2012; Awad et al., 2007; Stephens et al., 2010). This aligns with the notion that control processes are necessary for managing the complex task of generating meaningful, connected speech (Bourguignon et al., 2018; Bourguignon & Gracco, 2019; Geranmayeh et al., 2014, 2016; Wise & Geranmayeh, 2016). The importance of domain-general cognitive mechanisms for speech regulation has also been illustrated in neuropsychological studies (e.g., Barker et al., 2017; Marini, 2012; Marini et al., 2011). For example, Barker et al., (2017) found that mild stroke patients with right hemisphere damage exhibited significantly lower global coherence, and moreover, they exhibited attentional and executive impairments that were correlated with difficulties in producing coherent speech. The current study adds to this literature, underscoring the critical involvement of the MDN in orchestrating executive and attentional processes necessary for sustaining speech coherence, and further highlighting the joint roles of domain-general cognitive mechanisms and language-specific functions in speech production (Bourguignon & Gracco, 2019).

The effect of coherence in the SCN was weaker and not statistically significant. Previous neuropsychological work has demonstrated that patients with semantic aphasia, who exhibit deficits specific to semantic control, struggle to maintain global coherence, but can manage local coherence (Hoffman et al., 2020). Additionally, Hoffman et al., (2018) highlighted the association between *semantic selection* mechanisms - the ability to focus on specific semantic properties while avoiding dominant associations - and global coherence. In natural discourse production, a conversational cue may activate a wide of range of semantic information, not all of which is relevant to the conversational goal. Enhanced semantic selection mechanisms may allow the speaker to effectively select those aspects of knowledge that are pertinent to the conversation at hand. Supporting this, neuroimaging research in healthy older adults has linked increased coherence to activation in the LIFG - and more specifically, the pars triangularis (BA 45) - a key area of the semantic control network linked to semantic selection mechanisms (Hoffman, 2019). The weaker effect of coherence on SCN activation in the present study may stem from the fact that we controlled for more coherent speech being less semantically divergent. As we discuss later, the linking of divergent semantic concepts has been associated with increased SCN activity.

Next, we turn to the question of creativity in discourse. Our functional connectivity analyses highlighted increased interplay between the default and executive networks in the context of more divergent speech patterns. This finding suggests that the production of more semantically diverse speech is linked to interactions among a broad set of interconnected regions. Typically, the DMN is associated with internally directed cognition, such as daydreaming or mental simulation (Andrews-Hanna et al., 2018; AndrewslJHanna et al., 2014; Fox et al., 2015); while the MDN is linked to task-focused executive functions (Duncan, 2010; Fedorenko et al., 2013). Our results indicate that these networks may work in tandem when people generate more semantically diverse speech content. This aligns with numerous studies that have proposed that default and executive networks exhibit dynamic coupling during divergent thinking tasks, reflecting the complex interaction between associative and executive processing underlying creativity (e.g., Beaty, Benedek, et al., 2014; Beaty et al., 2015; Beaty, Silvia, et al., 2014). This default-executive interplay suggests that divergent thinking requires processes to aid the generation of novel ideas, as well as organisational control to manage and apply these ideas in a meaningful way. The results of the current study extend our understanding of the neural underpinnings of creativity, illustrating how these networks operate beyond controlled experimental tasks into the realm of spontaneous everyday speech.

Interestingly, we did not find that the Semantic Control Network was related to divergence in speech content. This is surprising because previous studies have implicated SCN regions, particularly the IFG, in the formation of novel links between concepts (Krieger-Redwood et al. (2023) and in divergent thinking more generally (CogdelllJBrooke et al., 2020). For example, Krieger-Redwood et al. (2023) found that the SCN is preferentially recruited when participants are required to generate unconventional links between items (*melon-bookcase*), while strong associations (*red-round*) are supported by automatic, associative patterns of retrieval underpinned by DMN activity. One potential reason for this discrepancy is that divergent responses in our study may have occurred when participants recalled idiosyncratic episodic memories, rather than deliberately forming novel semantic connections. For example, when asked *‘What sort of things usually happen at a wedding?’*, a participant might recall a particularly unusual wedding they attended that took place on a tropical island. While content generated in this way would be “semantically diverse”, it would not require the participant to generate new semantic connections, as they were simply retelling a familiar memory. Indeed, the DSI measure focuses on the semantic distance between ideas within the response but does not capture whether the content is novel to the individual themselves. The distinction between automatic and controlled retrieval is also crucial, as the SCN is typically engaged when the retrieval process requires more effort (e.g., Chiou et al., 2018; Gao et al., 2021; Whitney et al., 2011b; Zhang et al., 2021). Many of the prompts we used required participants to talk about familiar and well-established scenarios, which may have easily evoked episodic memories and familiar schemas, reducing the demand on the semantic control processes and thereby diminishing the degree to which the SCN was engaged.

Furthermore, Krieger-Redwood et al. (2023) emphasised the functional dissociation within the DMN. The ‘core’ DMN system (including regions such as AG, PCC, and ACC) is typically implicated in tasks that rely on strong semantic associations and episodic memory, such as autobiographical and self-referential thinking. In contrast, the dorsomedial subsystem (dmDMN; consisting of the ventral IFG, temporal and parietal regions) is engaged in the more effortful, controlled retrieval, where semantic control is required to navigate weaker links between concepts. This distinction underscores the flexibility of the DMN in supporting both automatic and controlled processing, depending on the complexity of the task. Crucially, the dmDMN shows overlap with the SCN, suggesting these networks are closely related. Future studies could investigate how varying the complexity and novelty of spontaneous speech tasks influences the interaction between the different DMN subsystems and control networks such as the MDN and SCN. Tasks designed to elicit more unconventional speech content may preferentially engage the dmDMN and SCN, reflecting the need for more controlled retrieval during creative idea generation.

In summary, our work explored the relationship between coherence and creativity in spontaneous speech, revealing a trade-off between maintaining focus on the original topic and introducing semantically diverse ideas. We demonstrated that speech coherence is supported by the Multiple Demand Network, which may facilitate the organisation and management of speech content; while creativity involves the interaction between the Default Mode and Multiple Demand Networks. These findings contribute to our understanding of how we balance structured communication with creative expression in everyday speech, and the neurocognitive networks that support these processes. Future work could further explore how task complexity influences the engagement of these networks.

## Acknowledgements

The research was supported by a BBSRC grant (BB/T004444/1). For the purpose of open access, the author has applied a Creative Commons Attribution (CC BY) licence to any Author Accepted Manuscript version arising from this submission.

