## Supplementary_Materials for "Balancing Act: A Neural Trade-off Between Coherence and Creativity in Spontaneous Speech"

**Description of Datasets**

***MD Dataset.*** This sample consisted of 30 younger adults (*M*_Age_= 20.9, range = 19-26), recruited through the undergraduate participant pool and compensated with course credit, and 23 older adults (*M*_Age_= 73.2, range = 65-89) recruited through the department’s volunteer panel. The younger sample had an average of 16.3 years of education, while the older sample had 15.0 years of education. These are unpublished data from student projects supervised by PH.

***18-ELife Dataset.*** This sample consisted of 30 younger adults (*M*_Age_= 19.3, *SD*_Age_ = 2.2, range = 18-30), recruited through the Psychology Department’s undergraduate participant pool, and 30 older adults (*M*_Age_= 76.0, *SD*_Age_ = 8.3, range = 61-91), recruited through the Department’s volunteer panel. The younger group had an average of 13.8 years of education, while the older group had 14.3 years of education. Previous analysis of these data, examining speech coherence in relation to executive and semantic abilities, has been published (see Hoffman et al., 2018).

***19-NC Dataset.*** The sample consisted of 15 older adults (*M*_Age_= 77.5, range = 67-92), who had an average of 15.0 years of education, recruited via the Department’s volunteer panel. These data have previously been used in a study examining the neural correlates of coherence in speech production (Hoffman, 2019).

***SL19 Dataset.*** The sample consisted of 25 young adult participants (*M*_Age_= 24, *SD*_Age_ = 4.4, range = 18-35, 21*F*, 4*M*) recruited from the student population. These data has been part of several prior analyses examining neural networks related to coherence (Morales et al., 2022), the neural correlates related to psycholinguistic properties of naturalistic discourse (Wu et al., 2022) and the neural encoding of meaning during speech production and comprehension (Patel et al., 2023).

##### **Table S1:** Demographic summary of datasets.

|  |  | Age | | | | f/m | No of Prompts |
| --- | --- | --- | --- | --- | --- | --- | --- |
| Group | *n* | *M* | *SD* | *Min* | *Max* | *Count* | *Count* |
| 18-MD | 30 | 20.9 | 1.41 | 19 | 26 | 22/8 | 12 |
| 19-MD | 23 | 72.26 | 5.96 | 65 | 89 | 13/10 | 12 |
| 18-EL | 60 | 47.53 | 29 | 17.98 | 91.41 | 37/23 | 14 |
| 19-NC | 15 | 76.91 | 8.09 | 65.81 | 91.41 | 9/6 | 20 |
| 19-SL | 25 | 23.83 | 4.29 | 18 | 35 | 21/4 | 12 |
| Combined | 153 | 47.14 | 27.83 | 17.98 | 91.41 | 102/51 | 27 |
| Young Adults | 85 | 21.08 | 3.32 | 17.98 | 35 | 65/20 | 26 |
| Older Adults | 68 | 75.53 | 7.75 | 61.65 | 91.41 | 37/31 | 20 |

**Study Design and Procedure**

***MD Dataset.*** Participants were presented with a prompt on a computer screen, which was followed by a tone that cued them to start speaking about the topic. They were asked to stop speaking when they heard the second tone. Participants responded to prompts in three conditions, each with a different set of goals. In the “Interesting” condition, participants were instructed to make their responses as *interesting, fascinating, and entertaining* as possible; in the “Clear” condition, participants had to keep their responses as *clear, focused, and simple* as possible. Finally, participants were given no instructions in the “Baseline” condition. The order of conditions was counterbalanced between participants. Each participant responded to 12 prompts.

***18-ELife Dataset.*** In this study, speech was elicited under two conditions, dual-task and speech-only, with 7 speech samples elicited in each condition. On speech-only trials, participants were presented with a written prompt and asked to press a key when they were ready to begin. They were then asked to speak about the topic until they heard a tone, with each trial lasting 60s. The dual-task trials, where participants produced speech while completing an attention demanding task, were excluded from the current analysis.

***19-NC Dataset.*** Participants were presented with a series of prompts addressing a range of semantic knowledge while undergoing fMRI. At the start of each trial, there was a preparation phase, where the written prompt was presented on screen for 8s. When they saw a green circle, participants were asked to speak about the subject for 50s, until the circle was replaced by a red cross. In the baseline trials, which lasted 15s, participants were asked to recite the nursery rhyme ‘Humpty Dumpty’. This baseline task was designed to invoke grammatically correct continuous speech without the need for generation of novel utterances. Each participant responded to 20 prompts.

***SL19 Dataset.*** In this study, participants were asked to produce and comprehend passages of naturalistic speech while being scanned. They were presented with two runs of the production and comprehension conditions in an alternating sequence, and the order of the runs was counterbalanced across participants. Each run lasted approximately 8 minutes and consisted of 6 speech trials and 5 baseline trials, randomly presented to each participant. The duration of the baseline trials was 10s, wherein participants recited or listened to the Humpty Dumpty nursery rhyme. For the purposes of this analysis, we focused on the speech production condition. First, in the preparation phase, participants saw a written prompt on screen for 6s. Then, they were asked to start speaking at the onset of a green circle, and to stop when this was replaced with a red cross. Each trial lasted 50s and participants were asked to try to stay on topic as far as they could. Prior to the experimental trials, participants were given practice trials to ensure they understood the instructions. Each participant responded to 12 prompts.

##### *Table S2: Participant-level correlations between measures in each individual sample and in the combined dataset.*

|  | Coherence - DSI | | Coherence – Words | | Coherence – Words | |
| --- | --- | --- | --- | --- | --- | --- |
| Group | *r* | *p* | *r* | *p* | *r* | *p* |
| MD | -.34 | **.01** | -.28 | **.04** | .46 | **<.001** |
| 18-EL | -.68 | **<.001** | -.17 | .18 | -.12 | .36 |
| 19-NC | -.32 | .23 | .34 | .20 | -.03 | .91 |
| 19-SL | -.26 | .19 | -.27 | .18 | .29 | .16 |
| Overall | -.52 | **<.001** | -.28 | **<.001** | .21 | **<.001** |

*Table S3: Demographic summary and list of prompts used across datasets.*

| Code | Participants | Datasets | Prompt |
| --- | --- | --- | --- |
| Ages | 127 | MD, EL, NC | What would it have been like to live in the Middle Ages? |
| Antarctica | 126 | MD, EL, NC | What would it be like to live in Antarctica? |
| Crime | 125 | MD, EL, NC | What do the police do when a crime has been committed? |
| Dog | 127 | MD, EL, NC | What sort of things do you have to do to look after a dog? |
| Mars | 127 | MD, EL, NC | Do you think it's a good idea to send people to live on Mars? |
| Morning | 125 | MD, EL, NC | What do people usually do when getting ready for work in the morning? |
| Restaurant | 126 | MD, EL, NC | Describe a typical visit to a restaurant. |
| Scotland | 127 | MD, EL, NC | Describe why someone might travel to Scotland on holiday |
| Season | 125 | MD, EL, NC | Which is your favourite season and why? |
| Storm | 125 | MD, EL, NC | What happens when a storm is forecast in the UK? |
| Train | 126 | MD, EL, NC | Describe the steps you would need to take if going somewhere by train. |
| University | 128 | MD, EL, NC | What are the advantages and disadvantages of going to university? |
| Christmas | 72 | EL, NC | What do people usually do on Christmas Day in the UK? |
| Queen | 75 | EL, NC | What sort of things does the Queen do on a typical day? |
| Election | 15 | NC | What happens during a general election in the UK? |
| Beverage | 40 | NC, SL | Describe how you would make a cup of tea or coffee |
| Wedding | 40 | NC, SL | What sort of things usually happen at a wedding? |
| Climate | 40 | NC, SL | Why are some people concerned about climate change? |
| Internet | 39 | NC, SL | Do you think the internet has improved people's lives? |
| Holiday | 39 | NC, SL | How would you prepare to go on holiday? |
| Diet | 25 | SL | Why is it important to have a balanced diet? |
| Food | 25 | SL | Describe the steps you’d take to order-in food. |
| Grocery | 25 | SL | Describe a typical visit to a grocery store. |
| Interview | 25 | SL | What would you recommend doing during a job interview? |
| New Year | 25 | SL | What do people usually do on New Year’s Eve in the UK? |
| Stress | 25 | SL | What sorts of things do people do to cope with stress? |
| Teacher | 24 | SL | What sort of things does a teacher do when at work? |

##### *Table S3: Participant-level correlations between measures in each individual sample and in the combined dataset.*

|  | Gc - DSI | | Gc – Words | | DSI – Words | |
| --- | --- | --- | --- | --- | --- | --- |
| Group | *r* | *p* | *r* | *p* | *r* | *p* |
| MD | -.34 | **.01** | -.28 | **.04** | .46 | **<.001** |
| 18-EL | -.68 | **<.001** | -.17 | .18 | -.12 | .36 |
| 19-NC | -.32 | .23 | .34 | .20 | -.03 | .91 |
| 19-SL | -.26 | .19 | -.27 | .18 | .29 | .16 |
| Overall | -.52 | **<.001** | -.28 | **<.001** | .21 | **<.001** |

*Figure S1: Unthresholded maps (beta values) of the effects of the regressors on neural activation. (a) Coherence effects; (b) DSI effects; (c) No of words effects*

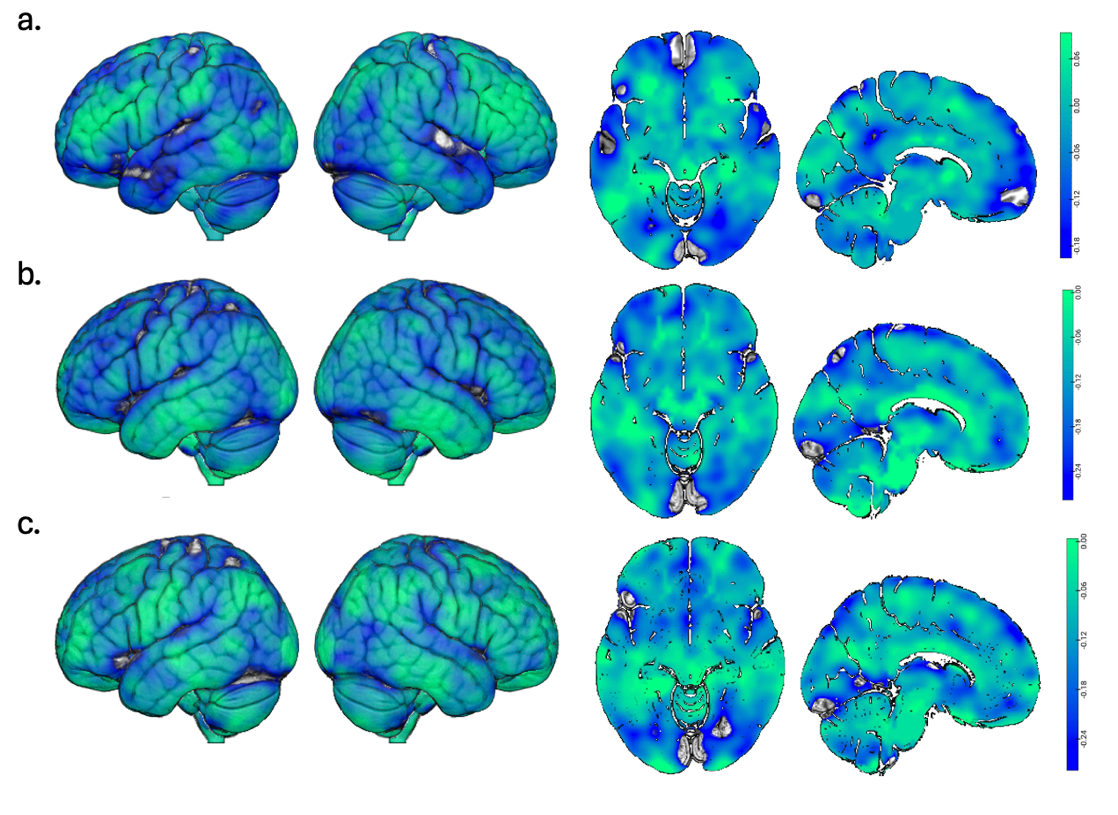
